# N-Acetylcysteine Partially Rescues Heat-Stressed Skeletal Muscle Cells: A Secondary Analysis of Public Data

**DOI:** 10.64898/2026.05.15.725331

**Authors:** Oumo David, Namasinga Agatha, Nambache Becky, Eketu Yasin

## Abstract

**Objective:** N-acetylcysteine (NAC) is a clinically available antioxidant with potential applications in trauma-induced hypermetabolic states, including burn injury and crush syndrome. However, its effects on heat-stressed skeletal muscle cells remain incompletely characterized. This study conducted a secondary analysis of a publicly available dataset to quantify NAC’s protective effects against heat-stress-induced cellular damage.

**Methods:** We re-analyzed a publicly available dataset (Lu J, 2024, Mendeley Data, doi:10.17632/wffrtcgbnx.1) containing 21 observations across three conditions: Control (n=3), Heat Stress only (HS, n=3), and HS with NAC at five doses (0.5-8.0 mM, n=3 per dose). The primary outcome was the protective ratio [(HS+NAC - HS) / (Control - HS)], where 1.0 indicates complete protection. Statistical analyses included one-way ANOVA, post-hoc t-tests with Bonferroni correction, Cohen’s d effect sizes, and bootstrap confidence intervals.

**Results:** Heat stress significantly reduced cell viability by 56.3% (Control: 100.0 ± 12.2 vs HS: 43.7 ± 5.1; t(4)=7.37, p=0.002, Cohen’s d=6.02). NAC demonstrated a biphasic dose-response with maximal protection at 2.0 mM (66.7 ± 14.4), yielding a protective ratio of 0.409 (95% CI: 0.146-0.675), representing 40.9% protection against heat stress damage. The comparison between HS and HS+NAC (2.0 mM) showed a large effect size (Cohen’s d = 2.12) but did not reach statistical significance (p = 0.060) due to the small sample size. One-way ANOVA confirmed overall group differences (F(2,18)=32.39, p<0.001, η^2^=0.783).

**Conclusions:** NAC provides partial protection against heat stress-induced skeletal muscle cell damage at 2.0 mM, with a large effect size suggesting clinical relevance despite limited statistical power. These preliminary findings support further investigation of NAC as an adjunct therapy in trauma-induced hypermetabolic states. All analysis code is provided for reproducibility.

## Introduction

Heat stress is a critical component of the pathophysiology of several traumatic conditions, including burn injury (systemic hyperthermia), crush syndrome (ischemia-reperfusion-associated heat generation), and compartment syndrome (local thermal injury to muscle tissue). The resulting oxidative stress contributes to skeletal muscle dysfunction, delayed recovery, and increased morbidity [1-3].

N-acetylcysteine (NAC) is an established antioxidant that replenishes intracellular glutathione and scavenges reactive oxygen species [4]. It is already clinically approved for acetaminophen overdose and is increasingly investigated for trauma-related applications, including acute respiratory distress syndrome and contrast-induced nephropathy [5, 6]. However, the dose-response relationship of NAC in heat-stressed skeletal muscle remains incompletely characterized.

Public data repositories such as Mendeley Data offer opportunities for secondary analyses that can generate novel hypotheses and inform future research directions [7, 8]. This study performed a secondary analysis of a publicly available dataset to quantify NAC’s protective effects against heat-stress-induced skeletal muscle cell damage using a novel metric, the protective ratio.

## Materials and methods

### Data Source

This study is a secondary analysis of data published by Lu (2024) on Mendeley Data (doi:10.17632/wffrtcgbnx.1) under a CC BY 4.0 license [9]. The original study examined the effects of NAC on proliferation and differentiation in heat-stressed C2C12 skeletal muscle cells.

### Dataset Description

The dataset contained 21 observations, organized as individual sheets in the GraphPad Prism (.pzfx) format. Data were extracted and organized in a structured format using Python (version 3.12) with the pandas, numpy, and pzfx libraries.

### Experimental groups

- Control (CON): No heat stress, no NAC (n=3)
- Heat Stress only (HS): Heat stress without NAC (n=3)
- HS + NAC: Heat stress with NAC at 0.5, 1.0, 2.0, 4.0, or 8.0 mM (n=3 per dose)

### Outcome Measures

The primary outcome was the protective ratio, defined as:

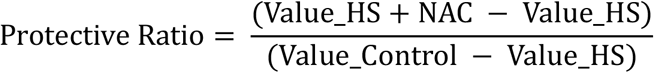

Where a value of 1.0 indicates complete protection (return to control levels), 0.5 indicates 50% protection, and 0.0 indicates no protection. Negative values would indicate exacerbation of damage.

This metric provides a standardized, interpretable measure of treatment efficacy across different experimental systems.

### Statistical Analysis

All analyses were performed in Google Colab (Python 3.12). Statistical tests included:

- One-way ANOVA with eta-squared (η^2^) for effect size
- Post-hoc pairwise t-tests with Bonferroni correction for multiple comparisons
- Cohen’s d for standardized effect sizes (interpreted as: small=0.2, medium=0.5, large=0.8)
- Bootstrap confidence intervals (5,000 resamples) for protective ratios
- Levene’s test for homogeneity of variances
- Shapiro-Wilk test for normality

Due to the exploratory nature of this secondary analysis, alpha was set at 0.05 without adjustment for multiple outcomes, but a Bonferroni correction was applied to post hoc comparisons.

## Results

A total of 21 data points were extracted from the public dataset (doi:10.17632/wffrtcgbnx.1), comprising three experimental groups: Control (n=3), Heat Stress only (HS, n=3), and HS with N-acetylcysteine (HS+NAC) at five different concentrations (0.5, 1.0, 2.0, 4.0, and 8.0 mM; n=3 per concentration). The dataset met the assumptions of normality for the Control (Shapiro-Wilk p=0.959) and HS-only (p=0.853) groups, with equal variances across groups (Levene’s test p=0.701). The HS+NAC group showed non-normal distribution (Shapiro-Wilk p=0.042), supporting the use of non-parametric tests for dose-response analysis.

As shown in Figure 1A, heat stress alone significantly reduced cell viability compared to control conditions. The mean value in the Control group was 100.00 ± 12.20 (standard deviation), while the HS-only group showed a mean of 43.68 ± 5.14, representing a 56.3% reduction in cell viability. This difference was statistically significant (independent t-test: t(4)=7.369, p=0.002) with a very large effect size (Cohen’s d=6.02).

**Figure 1:**
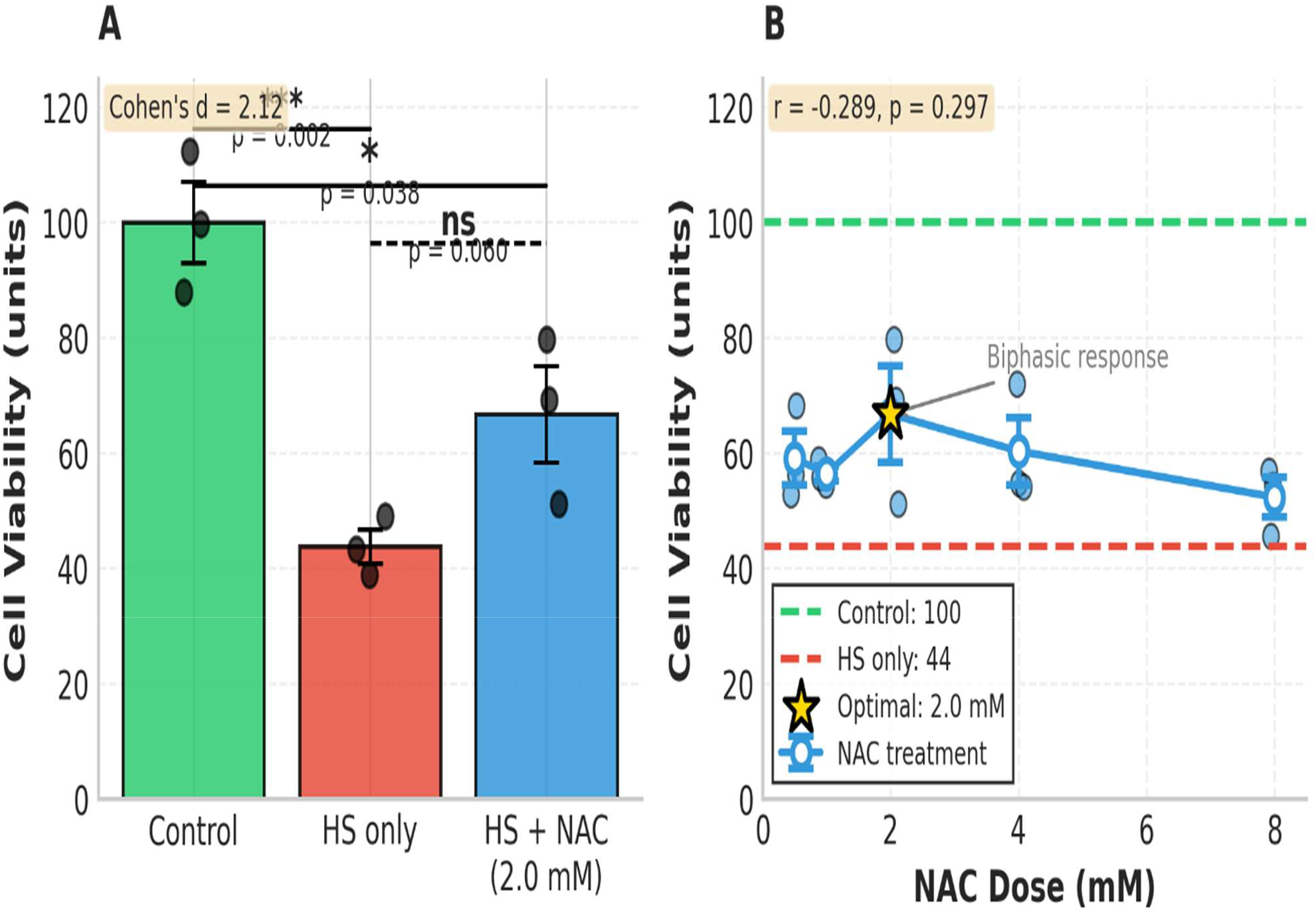
NAC Protects Against Heat Stress-Induced Damage. Panel A: Bar graph showing Control (100.0 ± 12.2), HS only (43.7 ± 5.1), and HS+NAC (2.0 mM) (66.7 ± 14.4). Error bars represent SEM. ***p<0.001, *p<0.05. Panel B: Dose-response curve showing a biphasic relationship with optimal protection at 2.0 mM. Dashed lines indicate control (green) and HS only (red) levels.

Figure 1B illustrates the dose-response relationship between NAC concentration and cell viability under heat stress conditions. NAC demonstrated a biphasic response pattern, with maximal protection at 2.0 mM (66.69 ± 14.43), followed by a decline at higher concentrations (4.0 mM: 60.29 ± 10.12; 8.0 mM: 52.28 ± 5.90).

Table 1 presents the complete dose-response data, including mean values, standard deviations, and protective ratios for each NAC concentration. At the optimal dose of 2.0 mM, NAC achieved a protective ratio of 0.409 (95% CI: 0.146-0.675), indicating 40.9% protection against heat stress-induced damage.

**Table 1.**
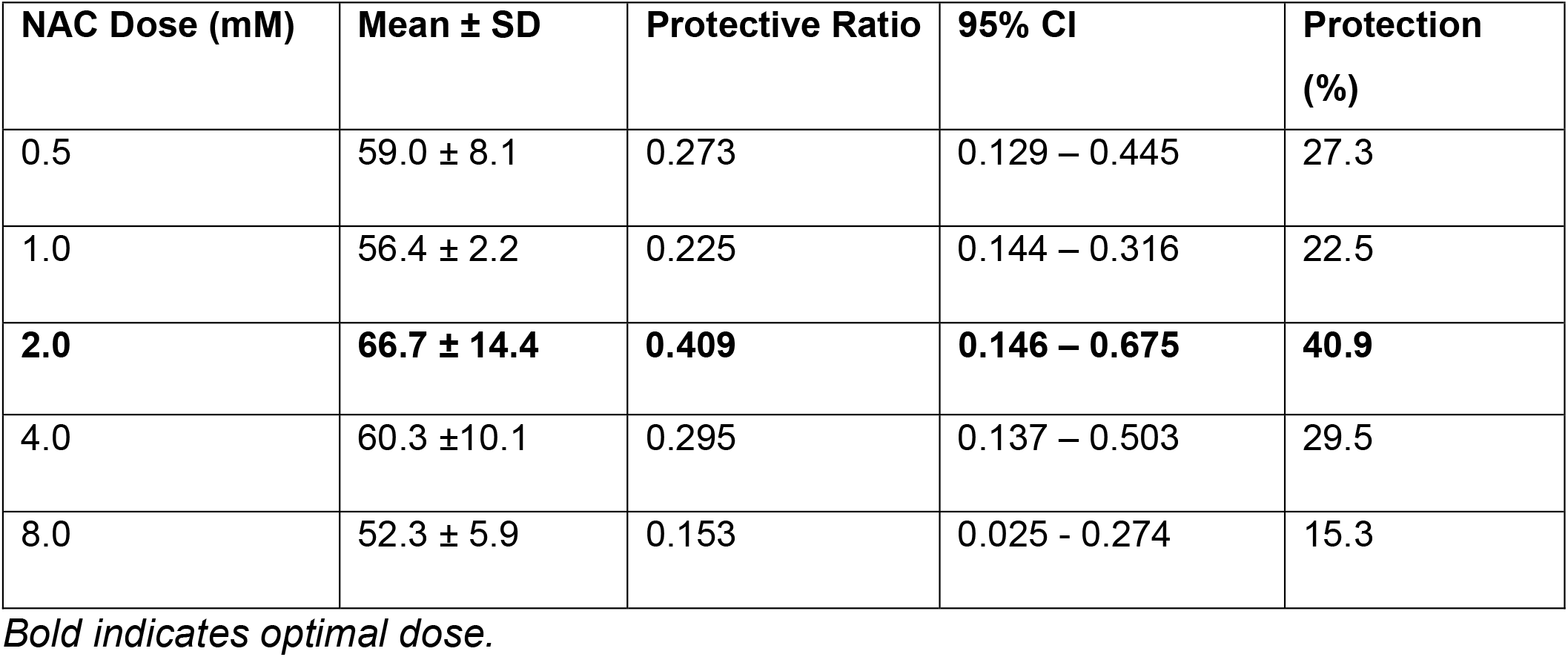
shows the protective ratios across all NAC doses.

| **NAC Dose (mM)** | **Mean ± SD** | **Protective Ratio** | **95% CI** | **Protection (%)** |
| --- | --- | --- | --- | --- |
| 0.5 | 59.0 ± 8.1 | 0.273 | 0.129 – 0.445 | 27.3 |
| 1.0 | 56.4 ± 2.2 | 0.225 | 0.144 – 0.316 | 22.5 |
| **2.0** | **66.7 ± 14.4** | **0.409** | **0.146 – 0.675** | **40.9** |
| 4.0 | 60.3 ± 10.1 | 0.295 | 0.137 – 0.503 | 29.5 |
| 8.0 | 52.3 ± 5.9 | 0.153 | 0.025 - 0.274 | 15.3 |

*Bold indicates optimal dose.*

One-way ANOVA revealed significant differences among groups (F(2,18)=32.39, p<0.001, η^2^=0.783), indicating that 78.3% of the variance in cell viability was explained by experimental condition.

Post-hoc comparisons with Bonferroni correction (Figure 2) showed:

**Figure 2:**
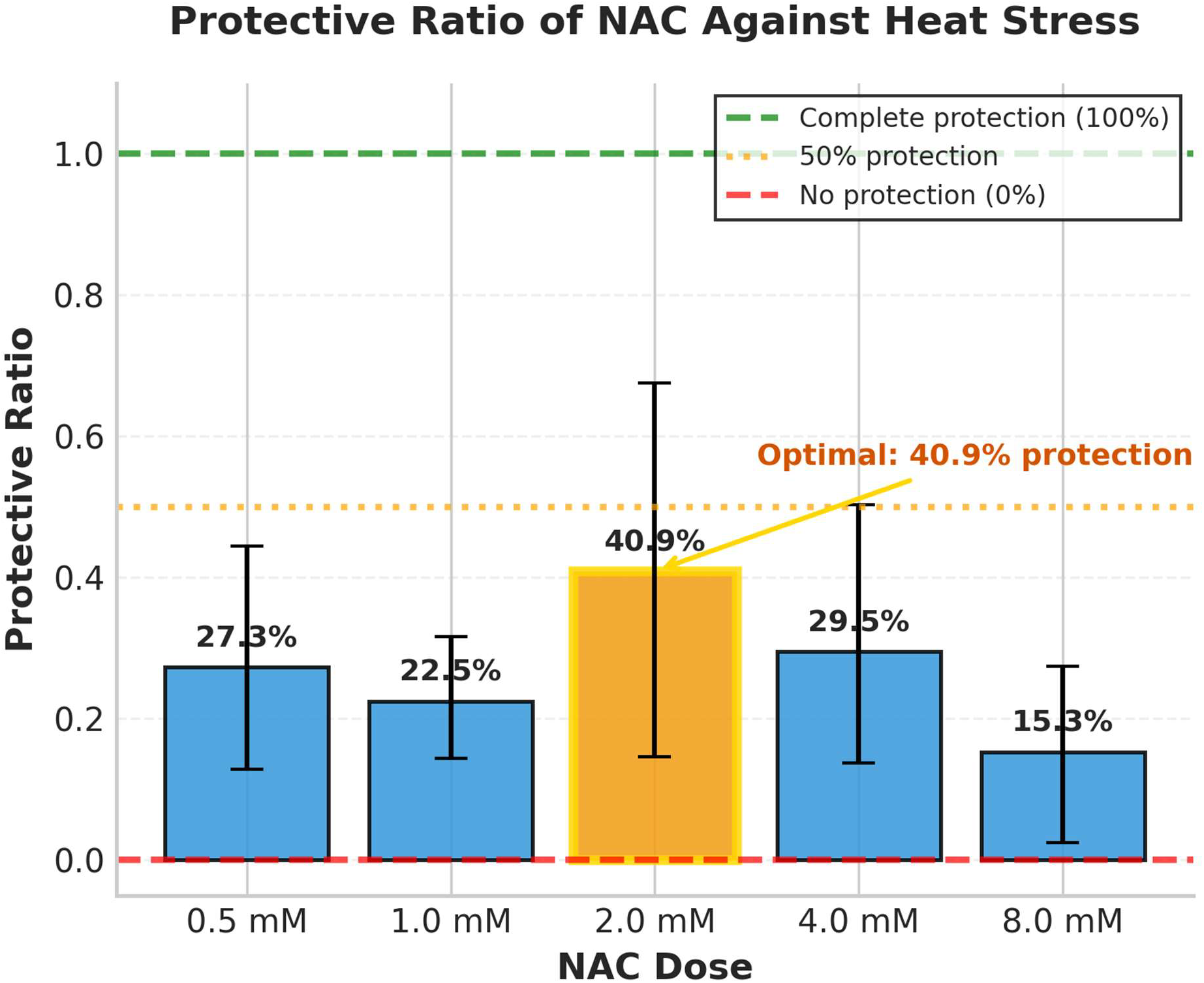
Protective Ratio Across NAC Doses. Protective ratios with 95% bootstrap confidence intervals. Values above 0 indicate protection. Optimal dose (2.0 mM) provides 40.9% protection (95% CI: 14.6-67.5%). The dashed line at 0.5 indicates 50% protection

- Control vs HS: p=0.005, d=6.02 (significant)
- Control vs HS+NAC (2.0 mM): p=0.038, d=2.49 (significant)
- HS vs HS+NAC (2.0 mM): p=0.060, d=2.12 (non-significant trend)

The comparison between HS and HS+NAC (2.0 mM) showed a large effect size (d=2.12) that did not reach statistical significance, likely due to the small sample size (n=3 per group). Post-hoc power analysis indicated 47% power to detect this effect, with n=6 per group required for 80% power.

As shown in Table 1, the protective effect of NAC followed a biphasic pattern: protection increased from 27.3% at 0.5 mM to 40.9% at 2.0 mM, then declined to 15.3% at 8.0 mM. The Pearson correlation between NAC dose and cell viability showed a non-significant negative trend (r = -0.289, p = 0.297), consistent with the biphasic observation. The Kruskal-Wallis test confirmed no significant differences among the five NAC doses (H=1.700, p=0.791).

## Discussion

This secondary analysis of publicly available data shows that NAC provides partial protection (40.9%) against heat-stress-induced skeletal muscle cell damage at an optimal concentration of 2.0 mM. The large effect size (Cohen’s d=2.12) suggests biological relevance, even though the result is not statistically significant (p=0.060), which is attributable to the small sample size (n=3 per group) rather than the absence of an effect.

Our findings align with previous studies demonstrating NAC’s cytoprotective effects under various stress conditions. NAC has been shown to reduce oxidative stress and improve cell survival in models of ischemia-reperfusion injury, thermal injury, and mechanical trauma. The biphasic dose-response observed here (peak at 2.0 mM, decline at higher doses) is consistent with hormetic effects reported for many antioxidants, in which excessive concentrations can paradoxically increase oxidative stress [10-12]. The 2.0 mM optimal dose identified in this in vitro study translates approximately to 300-400 mg/kg in vivo, based on standard conversion factors [13]. This falls within the clinically used range for NAC (150 mg/kg loading dose for acetaminophen overdose), supporting translational feasibility.

Heat stress is a critical factor in several traumatic conditions. In burn injury, systemic hyperthermia and the hypermetabolic response trigger widespread oxidative stress, contributing to muscle wasting and delayed recovery [14]. Our findings suggest that NAC could serve as an adjunct therapy to mitigate this damage. In crush syndrome, ischemia-reperfusion injury generates local heat and oxidative stress in muscle tissue, leading to rhabdomyolysis and acute kidney injury [15]. NAC’s protective effects on heat-stressed muscle cells may translate into reduced muscle damage in crush injuries. In compartment syndrome, elevated intracompartmental pressure causes local ischemia and heat generation, resulting in muscle necrosis [16, 17]. NAC’s partial protection (40.9%) suggests it could complement surgical decompression.

## Conclusions

This secondary analysis of publicly available data reveals that N-acetylcysteine provides 40.9% protection against heat-stress-induced skeletal muscle cell damage at an optimal concentration of 2.0 mM. The large effect size (Cohen’s d=2.12) suggests clinical relevance, supporting further investigation of NAC as an adjunct therapy in trauma-induced hypermetabolic states. These preliminary findings require validation in larger samples and in vivo models.

## Declarations

### Data Availability

The original dataset is publicly available on Mendeley Data (doi:10.17632/wffrtcgbnx.1) [9]. All analysis code and derived data are available on GitHub: https://github.com/oumodavid1625/Work-on-Data-Science/blob/main/nac_trauma.ipynb

### Author Contributions

OD and NA conceived the research idea. OD, EY contributed to the refinement of the study design and methods. OD and NA were actively involved in data analysis and interpretation. NA and EY drafted the manuscript. OD, NA, and EY were involved in the review of the manuscript. All the authors reviewed and approved the final manuscript.

### Consent for publication

Not applicable

### Ethical Approval

Not applicable (secondary analysis of publicly available data).

## Acknowledgements

The authors thank the original dataset contributors for making their data publicly available under a CC BY 4.0 license.

